# Soil respiration is correlated with rainfall and soil moisture at multiple temporal scales in a seasonal wet tropical forest

**DOI:** 10.1101/2024.01.03.574121

**Authors:** Dayani Chakravarthy, H. V. Raghavendra, Jayashree Ratnam, Mahesh Sankaran

## Abstract

Soil respiration is the second largest natural flux of carbon (C) between terrestrial ecosystems and the atmosphere, with tropical forests amongst the largest contributors of such soil-derived carbon efflux. With climate change expected to drive changes in both temperature and rainfall, our ability to predict responses of the carbon cycle in the future hinges upon an understanding of how these factors influence soil respiration (Rs). However, these relationships remain poorly characterised across the seasonal tropics, especially south Asia. Here, for two seasonal tropical sites in the Western Ghats of India, we characterised annual, seasonal and temporal variation in Rs and assessed its temperature and moisture sensitivity over 6 years. At both sites, Rs was positively correlated with temperature at the instantaneous scale, but showed no relationship with temperature at seasonal or annual scales. In contrast, Rs showed significant relationships with rainfall and soil moisture at all temporal scales. At the annual scale, respiration was negatively correlated with total annual rainfall. At the seasonal scale, wet season Rs was significantly lower than in the dry season. At the instantaneous scale, Rs showed a parabolic relationship with soil moisture, where soil respiration increased with soil moisture up to ∼21% and decreased beyond that point. Our results suggest that future responses of soil respiration in this seasonal tropical forest will largely be driven by changes in the Indian Summer Monsoon, not only mean annual precipitation, but also the frequency and intensity of extreme rainfall events.

## 1. Introduction

Tropical forests store about 55% of the global terrestrial carbon, of which nearly a third (∼32%) is sequestered in soils (Pan et al., 2011). Given that soils are so carbon rich, the release of carbon from soils is of much importance, and can play critical roles in regulating atmospheric CO_2_ concentrations and regional and global climate. Soil respiration (Rs) - comprising both autotrophic (root), and heterotrophic (bacterial and mycorrhizal) respiration - is a major contributor to the land-atmosphere carbon flux, releasing ∼75-98 Pg y^-1^ of C back to the atmosphere globally (Bond-Lamberty and Thomson, 2010a; Raich and Potter, 1995), with tropical forests amongst the largest contributors of these soil derived atmospheric carbon fluxes (Raich and Schlesinger, 1992; Warner et al., 2019).

Rates of soil respiration (Rs) are governed by complex interactions between a range of biotic and abiotic drivers, of which climatic factors such as temperature and moisture availability are particularly important (Bond-Lamberty and Thomson, 2010a; Hursh et al., 2017; Lei et al., 2021; Raich and Tufekciogul, 2000). Future projected changes in both temperature and rainfall patterns are thus likely to have significant impacts on soil derived land-atmosphere fluxes of CO_2_, with potential feedbacks on regional and global climates. Although temperature explains much of the variability in Rs across different biomes, other factors such as rainfall, soil carbon and carbon inputs to soils (e.g., litterfall) appear to be equally or more important in explaining observed variability in Rs within biomes (Bond-Lamberty and Thomson, 2010a; Huang et al., 2020; Hursh et al., 2017; Raich et al., 2002). For example, in tropical biomes, Rs is often uncorrelated or only weakly correlated with temperature (Davidson et al., 2000; Rubio and Detto, 2017), possibly a consequence of the low intra-annual variability in temperatures experienced by these biomes when compared to temperate ecosystems.

On the other hand, soil moisture has been shown to be an important driver of Rs in tropical systems (Adachi et al., 2009; Hashimoto et al., 2004; Rubio and Detto, 2017; Valentini et al., 2008; Wood et al., 2013). However, the exact nature of this relationship is unclear and appears variable, with studies reporting positive (Hashimoto et al., 2004), negative (Epron et al., 2006) and parabolic relationships of Rs with soil moisture (Adachi et al., 2009; Gaumont-Guay et al., 2006; Rubio and Detto, 2017; Sotta et al., 2004; Takahashi et al., 2011). Soil moisture can also mediate the effect of temperature on Rs, with low moisture typically constraining the response of Rs to temperature (Deng et al., 2012; Flanagan and Johnson, 2005; Hursh et al., 2017; Wen et al., 2006; Wood et al., 2013), although the opposite – higher temperature sensitivity of Rs at low soil moisture - has also been observed (Guntiñas et al., 2013).

As climate predictions for the future include changes in both temperature and rainfall patterns (IPCC, 2023), our ability to predict responses of the carbon cycle hinges upon a better understanding of the factors that influence soil carbon fluxes across different biomes. In particular, the limited numbers of empirical observations from tropical forests, which have amongst the highest predicted Rs rates, preclude good estimates of the variance in these rates, and have been identified as a major factor limiting robust global estimates of Rs, and for calibrating and reducing uncertainty in earth-system models (Hursh et al., 2017). The tropical forests of South Asia, in particular, are severely under-represented in most global syntheses of soil respiration (Akande et al., 2023; Bond-Lamberty and Thomson, 2010b, 2010a; Hursh et al., 2017; Lei et al., 2021).

In this study, we characterise the response of soil respiration to changes in moisture availability and temperature in two seasonal wet evergreen forest plots in the Western Ghats, India. These forests differ from other seasonal tropical wet forests in the length of the dry season (> 6 months) and in the intensity of rainfall they receive during the wet season (>2500 mm). Specifically, our objectives were to 1) quantify and characterise inter-annual and seasonal variation of soil respiration and 2) examine the effects of temperature and moisture availability on soil respiration rates at annual, seasonal and instantaneous temporal scales.

## 2. Methods

### 2.1 Site description

We conducted the study at two long-term wet evergreen forest monitoring sites – Hosagadde (14.478° N, 74.760° E) and Mulagunda (14.471°N,74.693 °E) – located in the Uttara Kannada district in the state of Karnataka in the central Western Ghats, India. Both sites are at an elevation of ∼550 m and have a long-term mean annual temperature of 24°C. The mean annual rainfall during the study period was 3730 mm and 4228 mm at Hosagadde and Mulagunda, respectively (University of East Anglia Climatic Research Unit; Harris et al., 2020). We assigned months with more than 100 mm rainfall as wet season (June – October), and months with <100 mm rainfall as the dry season (November – May). Soils in the region have been classified as a mixture of Eutric Nitosols and Acrisols under the FAO system (FAO-UNESCO, 1988; Krishnaswamy et al., 2017; Rome, FAO, 1998). From our measurements, soils in Hosagadde have 28.1% clay, 63.5 % sand and a water holding capacity of 83.22%, and soils in Mulagunda have 21.6% clay, 70.41% sand and a water holding capacity of 77.41%.

### 2.2 Field Measurements

We established 1ha permanent vegetation monitoring plots at Hosagadde and Mulagunda in 2013 and 2014, respectively, based on the RAINFOR-GEM (Marthews et al., 2014) and CTFS protocols (Condit, 1998). All trees >10 cm girth at breast height (GBH) in plots were identified, spatially mapped, tagged and their GBH measured. Plots were re-censused annually. We additionally quantified soil respiration in plots following the protocols outlined in Marthews et al., (2014). In each plot, we established 25 polyvinyl carbonate (PVC) soil collars (10 cm height and 10.6 cm diameter) by inserting them 5-6 cm into the soil. We used portable Infra-red gas analysers (IRGA) EGM-4 and EGM5 (PP Systems, USA) to measure instantaneous Rs (g CO_2_ m^-2.^h^-1^). Soil respiration was measured fortnightly from January 2014 to August 2020 in Hosagadde, and from January 2015 till August 2020 in Mulagunda.

Along with instantaneous collar-level Rs readings, we measured soil temperature using a digital thermometer with a 125mm stainless steel penetration probe (Hanna instruments). We also quantified instantaneous soil moisture at 30 cm depth during each sampling occasion using a FieldScout TDR 200 soil moisture meter (Spectrum Technologies, USA) at 3 points beside each soil collar, which were then averaged. For ambient air-temperature measurements, we used data from Thermochron i-button loggers (Maxim Integrated) placed at the centre and four corners of the 1 ha plot which recorded temperatures at 30 min intervals. Daily rainfall measurements were obtained from a tipping bucket rain gauge installed adjacent to the plot (Krishnaswamy et al., unpublished data).

### 2.3 Soil Respiration measurement and calculations

We calculated Rs following Marthews et al. (2014) as follows:

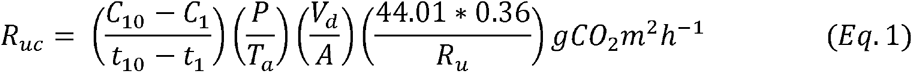

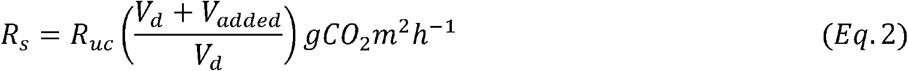

Here, *R*_*uc*_ is the uncorrected soil respiration (g CO2 m^-2^ h^-1^), estimated as the difference in CO_2_ concentrations C_*1*_ and *C*_*10*_, measured between times *t*_*1*_ and *t*_*10*_, where *t*_*10*_ was the last reading and *t*_*1*_ was typically 40 seconds before t_10_, *P* is the ambient pressure (mb) at time *t*_*10*_ as recorded by the IRGA, *T*_*a*_ is air temperature at time *t*_*10*_ in Kelvin, *V*_*d*_ the volume of the soil respiration chamber (0.001229 m^3^), *A* is the ground area of the soil collar over which R_uc_ was measured, and *R*_*u*_, the Universal Gas Constant (8.314 J K^-1^ mol^-1^). *R*_*s*_ is the soil respiration corrected for the additional volume of the soil collar *V*_*added*_ (in m^3^) based on the *h* cm of PVC collar protruding above the soil surface.

In order to maintain data-quality during field measurements, we compared the CO_2_ concentration at the end of each measurement to that at the start, and only included measurements where we observed a linear increase of CO_2_ between t_1_ and t_10._ (Marthews et al., 2014). We also repeated measurements where the end CO_2_ value was lower than the start value. If the end measurement was equal to the stat measurement, i.e., no respiration, we repeated the measurement for a total of three times before recording zero respiration. Additionally, during data processing, we excluded eight negative Rs values (4 each in Hosagadde and Mulagunda) that were missed during the field data check. Annual estimates of Rs at the plot level were obtained by spatially averaging hourly flux rates (gCO2.m^-2^.h^-1^) across all 25 collars in plots and scaling these up for the corresponding year (average hourly flux rates x 24 × 365). Seasonal fluxes were similarly obtained by first estimating average hourly flux rates for each season, and then scaling them up for the duration of the season.

### 2.4 Statistical methods

We assessed the effects of temperature and moisture availability on Rs at multiple temporal scales – annual, seasonal and instantaneous. At the annual scale, we used linear models to test the relationship between annual Rs and a) annual rainfall, b) mean annual temperature, c) number of rainy days, and d) number of days with heavy rainfall (>50mm).

At the seasonal scale, we used a linear mixed effects model (LMMs) to test whether Rs values differed between wet and dry seasons, with season (wet, dry) included as a fixed factor and soil collar number nested within year within site as a random factor to account for inherent differences between soil collars within each site, and non-independence of measurements over time at each collar. We had 11-24 data points/collar/year/site for a total of 6886 observations.

Finally, we used linear mixed effects models to test whether spatially averaged temperature and moisture levels could predict instantaneous Rs. For this, we used spatially averaged Rs (averaged over 25 collars) within each sampling period for each site as the response variable and spatially averaged temperature (as recorded by the IRGA) and a quadratic term with spatially averaged soil moisture and site ID as fixed factors. We used the calendar year as a random factor to account for inter-annual differences in soil respiration rates. We tested two candidate models: one which included an interaction term between temperature and soil moisture, and one without. In all, we had 13-24 sampling times/year/site and a total 276 spatially averaged Rs values. Rs values were square root transformed to meet model assumptions of linearity and homoscedasticity.

We used the R packages *glmmTMB* for all models, *DHARMa* for model diagnostics, *boxcox* to inform us about suitable transformations to meet model assumptions, *ggeffects, ggpubr* and *ggplot2* for data visualization and to plot model predictions, and *dplyr* for data cleaning and organisation. All analyses were carried out using R version 4.3.0 (R Core Team 2023).

## 3. Results

### 3.1. Rainfall, soil moisture and temperature seasonality

During the study period, annual rainfall in Hosagadde varied between 2800 - 5300 mm and in Mulagunda between 3700 - 7600 mm. Rainfall was very seasonal, occurring primarily during the south-west monsoon from June to October. Hosagadde received very heavy rains (>1000mm month^-1^) in July and August and Mulagunda between June and August. Outside of the monsoon, both sites received <100 mm rainfall month^-1^ (Fig 1A). Similar to rainfall variations, soil moisture levels were also strongly seasonal (Fig 1B). In both sites, they were lowest from February to April (average volumetric water content of 13.9% and 9.1%) and highest between July and October (35.6% and 30.2%). Mean annual temperatures during the study period at both sites were ∼22°C. Soil temperatures (at 125mm depth) ranged between 17-26°C in Hosagadde and 19 to 30°C in Mulagunda (Fig 1C).

**Figure 1:**
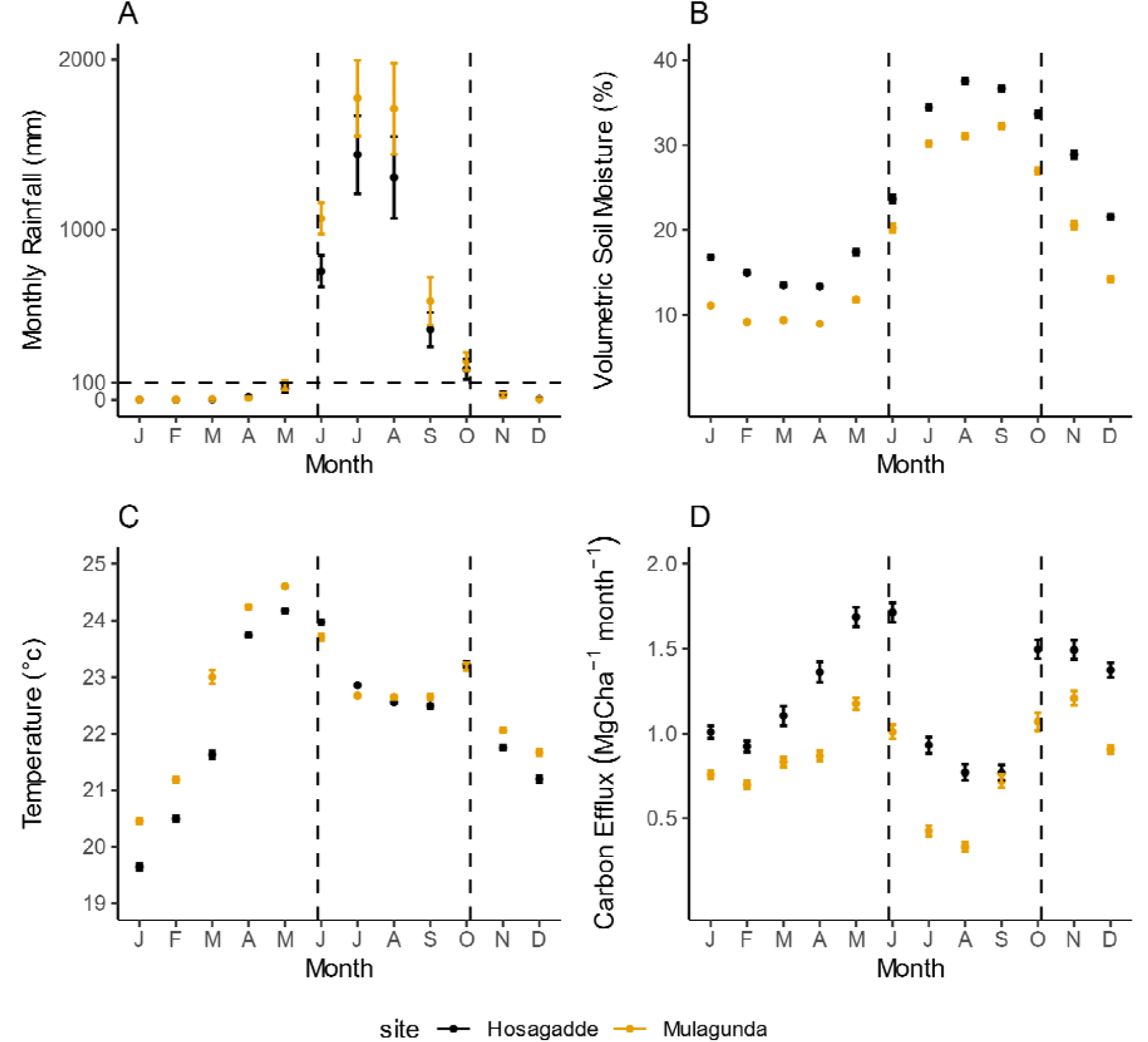
Monthly Variation in means and standard errors of A) Monthly Rainfall B) Soil Moisture C) Soil surface temperature and D) Soil Respiration (Mg C. ha^-1^. month^-1^). Months within dashed lines (Jun - Oct) represent the wet season.

### 3.2 Soil Respiration

Annual Rs ranged from 10.9 – 20.2 Mg C ha^-1^y^-1^ in Hosagadde (mean= 14.58, SD=3.38 cv=0.23), and 8.44 – 12.05 Mg ha^-1^ y^-1^ in Mulagunda (mean = 9.85, SD=1.6, cv=0.16) (Fig. S1) and was negatively related to total annual rainfall (p < 0.01) (Fig 2A). Annual Rs was not correlated with annual mean temperature (p=0.8), or number of rainy days in a year (p=0.23) (Table S1). However, it was significantly negatively correlated with the number of days where the rainfall was higher than 50 mm (p=0.002) (fig. 2B). Soil respiration in both sites was significantly lower during the wet season than during the dry season (p<0.001) (Fig 2C). In both sites, Rs was strongly seasonal, reaching high levels pre- and post-monsoon, and were lowest during the monsoon in July and August, months which received >1000mm rainfall (Fig 1D). At the instantaneous scale, the model without the interaction term between temperature and soil moisture had better support (lower AIC, Table S3) than the one with the interaction term included. Spatially averaged instantaneous Rs showed a parabolic relationship with soil moisture at both sites (significant quadratic term; p<0.001) with respiration peaking at ∼21% soil moisture (Fig 2D). For any given soil moisture level, respiration rates increased with increasing soil temperatures (Fig 2D).

**Figure 2:**
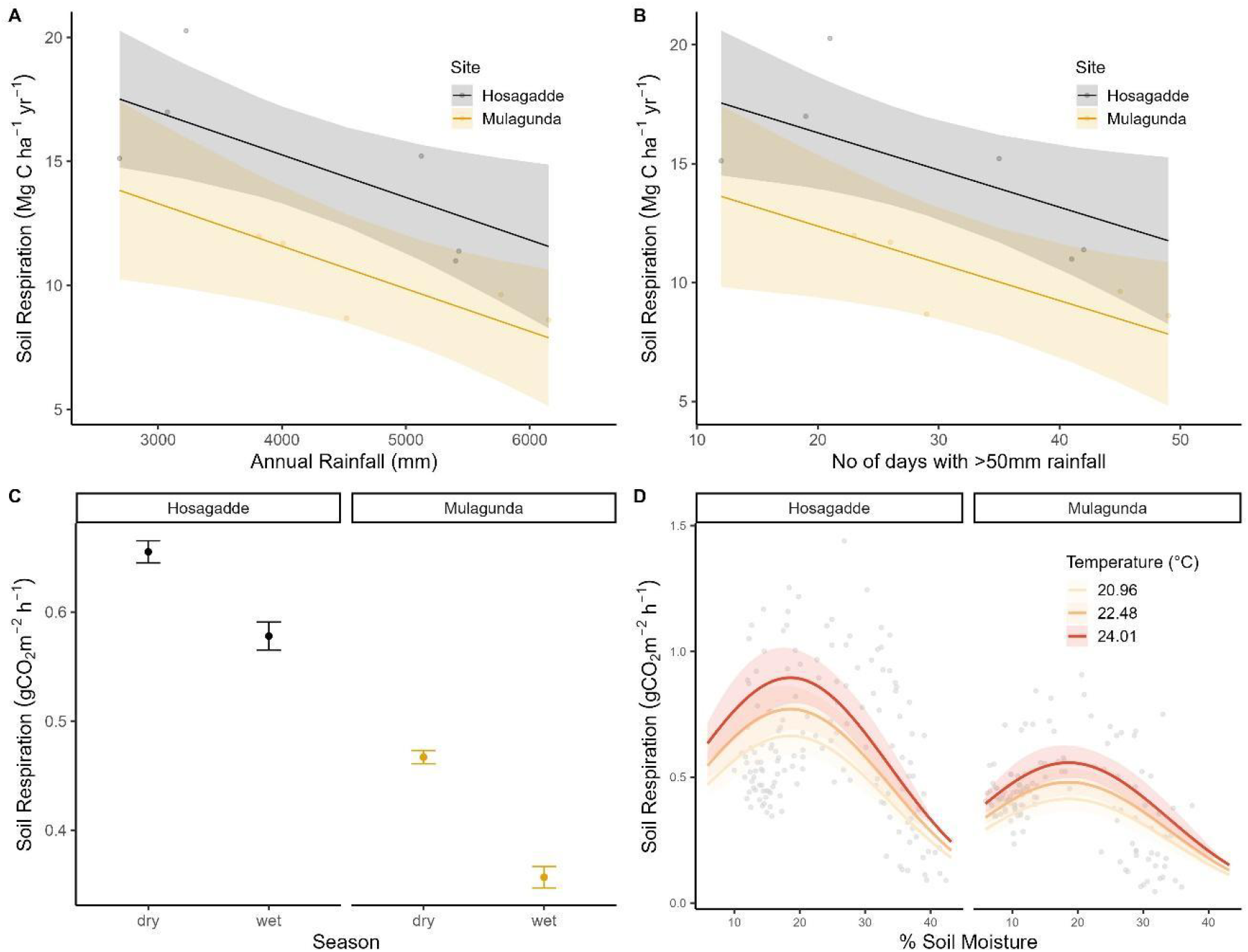
Soil respiration was correlated with rainfall and soil moisture at annual (A&B), seasonal (C) and instantaneous (D) timescales. In A, B &D, lines and bands show estimates and confidence intervals from models and points show data. In C, points and bars show means and standard errors of instantaneous soil respiration within each season.

## 4. Discussion

In this highly seasonal rainforest in the Western Ghats, rainfall was the dominant driver of soil respiration (Rs), influencing Rs at annual, seasonal and instantaneous time scales. Rs was lowest during periods of high soil moisture availability, i.e. during the wet-season in wetter years, and highest when soil moisture was lower, i.e. during the dry season and in drier years. In contrast, temperature was only of secondary importance, with warmer temperatures leading to higher rates of instantaneous CO_2_ efflux for any given soil moisture level.

Instantaneous soil respiration showed a parabolic relationship with soil moisture availability, peaking at ∼21% VWC and declining with both increases and decreases in soil moisture. Dry conditions can reduce Rs because of limited water availability for both microbial and root respiration, while high moisture availability can reduce rates of CO_2_ diffusion in waterlogged soils while also reducing microbial activity and root respiration as a result of soils becoming anoxic (Adachi et al., 2009; Cusack et al., 2019; Gaumont-Guay et al., 2006; Rubio and Detto, 2017; Valentini et al., 2008; Wood et al., 2013). Similar parabolic relationships of Rs with soil moisture have also been reported from other tropical forests across the globe (Adachi et al., 2009; Cusack et al., 2019; Koehler et al., 2009; Matson et al., 2017; Schwendenmann et al., 2003; Sotta et al., 2004). The soil moisture at peak respiration at our sites is comparable with reported values from Southeast Asian forests (∼21%) (Adachi et al., 2009; Takahashi et al., 2011), but lower than those from the Neotropics (30-40%; Sotta et al., 2004; Gaumont-Guay et al., 2006; Rubio and Detto, 2017; Wood et al., 2013), potentially a consequence of differences in soil texture across these sites. Soil water content for optimal Rs has previously been shown to be correlated with soil texture and clay content (Balogh et al., 2011; Cable et al., 2008).

Patterns of seasonal Rs in our sites are consistent with earlier results from neo-tropical forests that report lower Rs in wet seasons relative to dry seasons in high rainfall sites (>4000 mm MAP). In contrast, studies from drier sites report higher wet-season Rs relative to dry-season Rs (Cusack et al., 2019; Fernandes et al., 2002; Hashimoto et al., 2004; Koehler et al., 2009; Matson et al., 2017; Rubio and Detto, 2017; Schwendenmann et al., 2003; Sotta et al., 2006, 2004). Our results are also in accordance with experimental dry-down studies that report increases in Rs following rainfall reductions in wet sites, and decreases in Rs with drought in drier sites (Cleveland et al., 2010; van Straaten et al., 2011; Wood et al., 2013). Thus, at a global scale, the nature of the relationship between seasonal Rs and rainfall is likely determined by where sites lie along the dry-wet gradient.

In both our study sites, annual soil respiration was negatively correlated with the frequency of high rainfall events (>50mm day^-1^), but was not correlated with the frequency of rainy days, indicating that rainfall intensity is a key factor in limiting soil respiration in these forests. Not only do these sites receive extremely high rainfall, most of this rainfall occurs during the five monsoon months, with the wettest months receiving >1000mm rainfall month^-1^, while the rest of the year is mostly dry. Inhibition of soil respiration following extreme rain events has also been previously reported from other Asian monsoonal forests (Yu et al., 2021), and suggest that future increases in the frequency and intensity of extreme rain events are likely to significantly impact annual soil CO_2_ effluxes in these forests. Temperature, on the other hand, did not appear to be a strong driver of annual soil respiration rates in our study site, as has also been reported from other tropical forests (Deng et al., 2018; Rubio and Detto, 2017), possibly because it was largely invariant, with a range of less than 10 degrees in both of our sites.

Despite our two study sites being in close proximity to one another, soil CO_2_ efflux rates differed substantially between the two sites. Mean annual Rs ranged from 8.44 – 12.05 Mg.ha^-1^.yr^-1^ (mean = 9.85 Mg.ha^-1^.yr^-1^) at Mulagunda, and from 10.9 – 20.2 Mg.ha^-1^.yr^-1^ (mean = 14.58 Mg.ha^-1^.yr^-1^) at Hosagadde, as compared to the global mean of ∼13.1 Mg.ha^-1^.yr^-1^ reported for tropical broadleaved forests (Raich et al., 2002; Warner et al., 2019). The higher rainfall levels experienced in Mulgunda (Fig 1a) is a potential contributor to the lower fluxes at this site. However, even for the same level of soil moisture, instantaneous Rs tended to be lower in Mulgunda compared to Hosagadde (Fig 2d). While the reasons underlying these differences are currently unclear, it is likely that other factors including edaphic properties – which have been shown to exert significant controls on soil respiration rates, potentially more so than climatic factors (Haaf et al., 2021), soil nutrient availability, litter production and availability of substrate for microbes also contribute to the observed differences (Davidson et al., 2002; Yao et al., 2021).

Our study, the first to detail soil respiration at multiple temporal scales from a highly seasonal rainforest in the Western Ghats of India contributes to the growing body of literature on controls of soil respiration across the tropics. The combination of very high rainfall and extreme seasonality in these sites places them in a unique climatic niche globally, and adds valuable insights into our broader understanding of the potential relationships between rainfall regimes and soil respiration. Our key finding that both high annual rainfall and extremely rainy days suppress soil respiration indicates that future changes in the climatology of the monsoon in this region is likely to significantly influence carbon effluxes from these system in the future. Experimental studies that manipulate the intensity and frequency of heavy rainfall events can help to better quantify these patterns and thereby better parameterize earth system models. Further, separation of autotrophic and heterotrophic components of respiration can allow us to quantify the sensitivity of each component and better understand the biology underlying carbon fluxes in this ecosystem.

## Acknowledgements

We would like to thank Mahesha from Amminhalli for collecting data throughout the study with HVR. Thanks to Jagdish Krishnaswamy and FERAL for providing us with rainfall data, Balachandra Hedge for maintaining the weather station at Hosagadde, Sandeep Pulla, Anand MO and Deepak Barua for discussions during analysis, and BEER Lab folks at NCBS, especially Manaswi Raghurama and Mayank Kohli for their detailed suggestions on the manuscript. Finally, we would like to thank the Karnataka Forest Department for granting permission to conduct the study.

## References

Adachi, M., Ishida, A., Bunyavejchewin, S., Okuda, T., Koizumi, H., 2009. Spatial and temporal variation in soil respiration in a seasonally dry tropical forest, Thailand. Journal of Tropical Ecology 25, 531–539. 10.1017/s026646740999006x

Akande, O.J., Ma, Z., Huang, C., He, F., Chang, S.X., 2023. Meta-analysis shows forest soil CO(2) effluxes are dependent on the disturbance regime and biome type. Ecol Lett 26, 765–777. 10.1111/ele.14201

Balogh, J., Pintér, K., Fóti, Sz., Cserhalmi, D., Papp, M., Nagy, Z., 2011. Dependence of soil respiration on soil moisture, clay content, soil organic matter, and CO2 uptake in dry grasslands. Soil Biology and Biochemistry 43, 1006–1013. 10.1016/j.soilbio.2011.01.017

Bond-Lamberty, B., Thomson, A., 2010a. Temperature-associated increases in the global soil respiration record. Nature 464, 579–82. 10.1038/nature08930

Bond-Lamberty, B., Thomson, A., 2010b. A global database of soil respiration data. Biogeosciences 7, 1915–1926. 10.5194/bg-7-1915-2010

Cable, J.M., Ogle, K., Williams, D.G., Weltzin, J.F., Huxman, T.E., 2008. Soil Texture Drives Responses of Soil Respiration to Precipitation Pulses in the Sonoran Desert: Implications for Climate Change. Ecosystems 11, 961–979. 10.1007/s10021-008-9172-x

Cleveland, C.C., Wieder, W.R., Reed, S.C., Townsend, A.R., 2010. Experimental drought in a tropical rain forest increases soil carbon dioxide losses to the atmosphere. Ecology 91, 2313–23. 10.1890/09-1582.1

Condit, R., 1998. The CTFS and the Standardization of Methodology, in: Condit, R. (Ed.), Tropical Forest Census Plots: Methods and Results from Barro Colorado Island, Panama and a Comparison with Other Plots, Environmental Intelligence Unit. Springer, Berlin, Heidelberg, pp. 3–7. 10.1007/978-3-662-03664-8_1

Cusack, D.F., Ashdown, D., Dietterich, L.H., Neupane, A., Ciochina, M., Turner, B.L., 2019. Seasonal changes in soil respiration linked to soil moisture and phosphorus availability along a tropical rainfall gradient. Biogeochemistry 145, 235–254. 10.1007/s10533-019-00602-4

Davidson, E.A., Savage, K., Bolstad, P., Clark, D.A., Curtis, P.S., Ellsworth, D.S., Hanson, P.J., Law, B.E., Luo, Y., Pregitzer, K.S., Randolph, J.C., Zak, D., 2002. Belowground carbon allocation in forests estimated from litterfall and IRGA-based soil respiration measurements. Agricultural and Forest Meteorology 113, 39–51. 10.1016/s0168-1923(02)00101-6

Davidson, E.A., Verchot, L.V., Cattânio, J.H., Ackerman, I.L., Carvalho, J.E.M., 2000. Effects of Soil Water Content on Soil Respiration in Forests and Cattle Pastures of Eastern Amazonia. Biogeochemistry 48, 53–69. 10.1023/a:1006204113917

Deng, Q., Hui, D., Zhang, D., Zhou, G., Liu, J., Liu, S., Chu, G., Li, J., 2012. Effects of precipitation increase on soil respiration: a three-year field experiment in subtropical forests in China. PLoS One 7, e41493. 10.1371/journal.pone.0041493

Deng, Q., Zhang, D., Han, X., Chu, G., Zhang, Q., Hui, D., 2018. Changing rainfall frequency rather than drought rapidly alters annual soil respiration in a tropical forest. Soil Biology and Biochemistry 121, 8–15. 10.1016/j.soilbio.2018.02.023

Epron, D., Bosc, A., Bonal, D., Freycon, V., 2006. Spatial variation of soil respiration across a topographic gradient in a tropical rain forest in French Guiana. Journal of Tropical Ecology 22, 565–574. 10.1017/s0266467406003415

FAO-UNESCO, 1988. FAO-UNESCO soil map of the world. Revised legend. Paris.

Fernandes, S.A.P., Bernoux, M., Cerri, C.C., Feigl, B.J., Piccolo, M.C., 2002. Seasonal variation of soil chemical properties and CO2 and CH4 fluxes in unfertilized and P-fertilized pastures in an Ultisol of the Brazilian Amazon. Geoderma 107, 227–241. 10.1016/s0016-7061(01)00150-1

Flanagan, L.B., Johnson, B.G., 2005. Interacting effects of temperature, soil moisture and plant biomass production on ecosystem respiration in a northern temperate grassland. Agricultural and Forest Meteorology 130, 237–253. 10.1016/j.agrformet.2005.04.002

Gaumont-Guay, D., Black, T.A., Griffis, T.J., Barr, A.G., Jassal, R.S., Nesic, Z., 2006. Interpreting the dependence of soil respiration on soil temperature and water content in a boreal aspen stand. Agricultural and Forest Meteorology 140, 220–235. 10.1016/j.agrformet.2006.08.003

Guntiñas, M.E., Gil-Sotres F, Leirós, M.C., Trasar-Cepeda, C., 2013. Sensitivity of soil respiration to moisture and temperature. Journal of Soil Science and Plant Nutrition 13, 445–461. 10.4067/S0718-95162013005000035

Haaf, D., Six, J., Doetterl, S., 2021. Global patterns of geo-ecological controls on the response of soil respiration to warming. Nat. Clim. Chang. 11, 623–627. 10.1038/s41558-021-01068-9

Harris, I., Osborn, T.J., Jones, P., Lister, D., 2020. Version 4 of the CRU TS monthly highresolution gridded multivariate climate dataset. Sci Data 7, 109. 10.1038/s41597-020-0453-3

Hashimoto, S., Tanaka, N., Suzuki, M., Inoue, A., Takizawa, H., Kosaka, I., Tanaka, K., Tantasirin, C., Tangtham, N., 2004. Soil respiration and soil CO2 concentration in a tropical forest, Thailand. Journal of Forest Research 9, 75–79. 10.1007/s10310-003-0046-y

Huang, N., Wang, L., Song, X.P., Black, T.A., Jassal, R.S., Myneni, R.B., Wu, C.Y., Wang, L., Song, W.J., Ji, D.B., Yu, S.S., Niu, Z., 2020. Spatial and temporal variations in global soil respiration and their relationships with climate and land cover. Sci Adv 6. https://doi.org/ARTNeabb8508 10.1126/sciadv.abb8508

Hursh, A., Ballantyne, A., Cooper, L., Maneta, M., Kimball, J., Watts, J., 2017. The sensitivity of soil respiration to soil temperature, moisture, and carbon supply at the global scale. Glob Chang Biol 23, 2090–2103. 10.1111/gcb.13489

Intergovernmental Panel on Climate Change (IPCC) (Ed.), 2023. Summary for Policymakers, in: Climate Change 2022 – Impacts, Adaptation and Vulnerability: Working Group II Contribution to the Sixth Assessment Report of the Intergovernmental Panel on Climate Change. Cambridge University Press, Cambridge, pp. 3–34. 10.1017/9781009325844.001

Koehler, B., Corre, M.D., Veldkamp, E., Sueta, J.P., 2009. Chronic nitrogen addition causes a reduction in soil carbon dioxide efflux during the high stem-growth period in a tropical montane forest but no response from a tropical lowland forest on a decadal time scale. Biogeosciences 6, 2973–2983. 10.5194/bg-6-2973-2009

Krishnaswamy, J., Chappel, N., Bhalla, R., Sankaran, M., Vaidyanathan, S., Badiger, S., Varghese, S., 2017. Hydrologic and Carbon Services in the Western Ghats: Response of Forest and Agroecosystems to Extreme Rainfall Events (Technical Report submitted to Ministry of Earth Sciences, Government of India.).

Lei, J., Guo, X., Zeng, Y., Zhou, J., Gao, Q., Yang, Y., 2021. Temporal changes in global soil respiration since 1987. Nat Commun 12, 403. 10.1038/s41467-020-20616-z

Marthews, T.R., Riutta, T., Oliveras Menor, I., Urrutia, R., Moore, S., Metcalfe, D., Malhi, Y., Phillips, O., Huaraca Huasco, W., Ruiz Jaén, M., 2014. Measuring tropical forest carbon allocation and cycling: a RAINFOR-GEM field manual for intensive census plots (v3. 0). Manual, Global Ecosystems Monitoring network.

Matson, A.L., Corre, M.D., Langs, K., Veldkamp, E., 2017. Soil trace gas fluxes along orthogonal precipitation and soil fertility gradients in tropical lowland forests of Panama. Biogeosciences 14, 3509–3524. 10.5194/bg-14-3509-2017

Pan, Y., Birdsey, R.A., Fang, J., Houghton, R., Kauppi, P.E., Kurz, W.A., Phillips, O.L., Shvidenko, A., Lewis, S.L., Canadell, J.G., Ciais, P., Jackson, R.B., Pacala, S.W., McGuire, A.D., Piao, S., Rautiainen, A., Sitch, S., Hayes, D., 2011. A large and persistent carbon sink in the world’s forests. Science 333, 988–93. 10.1126/science.1201609

Raich, J.W., Potter, C.S., 1995. Global patterns of carbon dioxide emissions from soils. Global Biogeochemical Cycles 9, 23–36. 10.1029/94gb02723

Raich, J.W., Potter, C.S., Bhagawati, D., 2002. Interannual variability in global soil respiration, 1980-94. Global Change Biology 8, 800–812. 10.1046/j.1365-2486.2002.00511.x

Raich, J.W., Schlesinger, W.H., 1992. The global carbon dioxide flux in soil respiration and its relationship to vegetation and climate. Tellus B 44, 81–99. 10.1034/j.1600-0889.1992.t01-1-00001.x

Raich, J.W., Tufekciogul, A., 2000. Vegetation and Soil Respiration: Correlations and Controls. Biogeochemistry 48, 71–90. 10.1023/a:1006112000616

Rome, FAO-UN, 1998. World Reference Base for soil Resources. FAO.

Rubio, V.E., Detto, M., 2017. Spatiotemporal variability of soil respiration in a seasonal tropical forest. Ecol Evol 7, 7104–7116. 10.1002/ece3.3267

Schwendenmann, L., Veldkamp, E., Brenes, T., O’Brien, J.J., Mackensen, J., 2003. Spatial and temporal variation in soil CO2 efflux in an old-growth neotropical rain forest, La Selva, Costa Rica. Biogeochemistry 64, 111–128. 10.1023/a:1024941614919

Sotta, E.D., Meir, P., Malhi, Y., Donato nobre, A., Hodnett, M., Grace, J., 2004. Soil CO2 efflux in a tropical forest in the central Amazon. Global Change Biology 10, 601–617. 10.1111/j.1529-8817.2003.00761.x

Sotta, E.D., Veldkamp, E., Guimarães, B.R., Paixão, R.K., Ruivo, M.L.P., Almeida, S.S., 2006. Landscape and climatic controls on spatial and temporal variation in soil CO2 efflux in an Eastern Amazonian Rainforest, Caxiuanã, Brazil. Forest Ecology and Management 237, 57–64. 10.1016/j.foreco.2006.09.027

Takahashi, M., Hirai, K., Limtong, P., Leaungvutivirog, C., Panuthai, S., Suksawang, S., Anusontpornperm, S., Marod, D., 2011. Topographic variation in heterotrophic and autotrophic soil respiration in a tropical seasonal forest in Thailand. Soil Science and Plant Nutrition 57, 452–465. 10.1080/00380768.2011.589363

Valentini, C.M.A., Sanches, L., de Paula, S.R., Vourlitis, G.L., de Souza Nogueira, J., Pinto, O.B., de Almeida Lobo, F., 2008. Soil respiration and aboveground litter dynamics of a tropical transitional forest in northwest Mato Grosso, Brazil. Journal of Geophysical Research: Biogeosciences 113, n/a-n/a. 10.1029/2007jg000619

van Straaten, O., Veldkamp, E., Corre, M.D., 2011. Simulated drought reduces soil CO2 efflux and production in a tropical forest in Sulawesi, Indonesia. Ecosphere 2, art119. 10.1890/ES11-00079.1

Warner, D.L., BondlJLamberty, B., Jian, J., Stell, E., Vargas, R., 2019. Spatial Predictions and Associated Uncertainty of Annual Soil Respiration at the Global Scale. Global Biogeochemical Cycles 33, 1733–1745. 10.1029/2019gb006264

Wen, X.-F., Yu, G.-R., Sun, X.-M., Li, Q.-K., Liu, Y.-F., Zhang, L.-M., Ren, C.-Y., Fu, Y.-L., Li, Z.-Q., 2006. Soil moisture effect on the temperature dependence of ecosystem respiration in a subtropical Pinus plantation of southeastern China. Agricultural and Forest Meteorology 137, 166–175. 10.1016/j.agrformet.2006.02.005

Wood, T.E., Detto, M., Silver, W.L., 2013. Sensitivity of soil respiration to variability in soil moisture and temperature in a humid tropical forest. PLoS One 8, e80965. 10.1371/journal.pone.0080965

Yao, Y., Ciais, P., Viovy, N., Li, W., Cresto-Aleina, F., Yang, H., Joetzjer, E., Bond-Lamberty, B., 2021. A Data-Driven Global Soil Heterotrophic Respiration Dataset and the Drivers of Its Inter-Annual Variability. Global Biogeochemical Cycles 35, e2020GB006918. 10.1029/2020GB006918

Yu, J.-C., Chiang, P.-N., Lai, Y.-J., Tsai, M.-J., Wang, Y.-N., 2021. High Rainfall Inhibited Soil Respiration in an Asian Monsoon Forest in Taiwan. Forests 12, 239. 10.3390/f12020239

