## Supplemental Table 3 for "Soil respiration is correlated with rainfall and soil moisture at multiple temporal scales in a seasonal wet tropical forest"

*Table S 3: Summary of generalised linear mixed model with instantaneous soil respiration as the predictor, temperature, soil moisture^2^ and site (Hosagadde, Mulagunda) as fixed response variables. We used collar ID nested within the calendar year as random effects.*

*Model 1: Soil respiration +0.0001 ~ temperature + soil moisture ^2^ + site + (1|year/collar), family = Gamma, link=log*

*Model 2: Soil respiration +0.0001 ~ temperature * soil moisture ^2^ + site + (1|year/collar), family = Gamma, link=log*

| **Model** | **1. Temperature**  **+**  **Soil Moisture^2^** | | **2. Temperature**  *****  **Soil Moisture^2^** | |
| --- | --- | --- | --- | --- |
| Response | Spatially averaged Rs  (g CO_2_ m^-2^h^-1^) | | Spatially averaged Rs  (g CO_2_ m^-2^h^-1^) | |
| *Predictors* | *Estimates* | *p* | *Estimates* | *p* |
| (Intercept) | 0.0406 | **<0.001** | 0.0481 | 0.181 |
| mean temp | 1.1029 | **<0.001** | 1.0954 | 0.360 |
| mean moist [1st degree] | 1.0832 | **<0.001** | 1.1046 | 0.697 |
| mean moist [2nd degree] | 0.9978 | **<0.001** | 0.9961 | 0.551 |
| site [Mulagunda] | 0.6230 | **<0.001** | 0.6241 | **<0.001** |
| mean temp × mean moist [1st degree] |  |  | 0.9991 | 0.933 |
| mean temp × mean moist [2nd degree] |  |  | 1.0001 | 0.788 |
| **Random Effects** | | | | |
| σ^2^ | 0.16 | | 0.16 | |
| τ_00_ | 0.02 _year_ | | 0.02 _year_ | |
| ICC | 0.10 | | 0.11 | |
| N | 7 _year_ | | 7 _year_ | |
| Observations | 278 | | 278 | |
| Marginal R^2^ / Conditional R^2^ | 0.383 / 0.446 | | 0.384 / 0.449 | |
| AIC | -105.532 | | -102.169 | |
