## Supplemental Table 2 for "Soil respiration is correlated with rainfall and soil moisture at multiple temporal scales in a seasonal wet tropical forest"

*Table S 2: Summary of generalised linear mixed model with instantaneous soil respiration as the predictor, site (Hosagadde, Mulagunda) and season (wet, dry) as fixed response variables. We used collar ID nested within the calendar year, nested within site ID as random effects.*

*Formula:*

*Soil respiration+0.0001 ~ season + site + (1|site/year/collar), family = Gamma, link=log.*

| Response | Instantaneous Rs  (g CO_2_ m^-2^h^-1^) | | |
| --- | --- | --- | --- |
| *Predictors* | *Estimates* | | *p* |
| (Intercept) | 0.6235 | **<0.001** | |
| season [wet] | 0.8041 | **<0.001** | |
| site [Mulagunda] | 0.6831 | **0.001** | |
| **Random Effects** | | | |
| σ^2^ | 0.50 | | |
| τ_00_ _collar:year:site_ | 0.09 | | |
| τ_00_ _year:site_ | 0.04 | | |
| τ_00_ _site_ | 0.00 | | |
| N _collar_ | 25 | | |
| N _year_ | 7 | | |
| N _site_ | 2 | | |
| Observations | 6936 | | |
| Marginal R^2^ / Conditional R^2^ | 0.087 / NA | | |
