## Supplementary figures and images for "Soil respiration is correlated with rainfall and soil moisture at multiple temporal scales in a seasonal wet tropical forest"

### Supplemental Figure 1

*Figure S 1: Annual Soil respiration estimates*

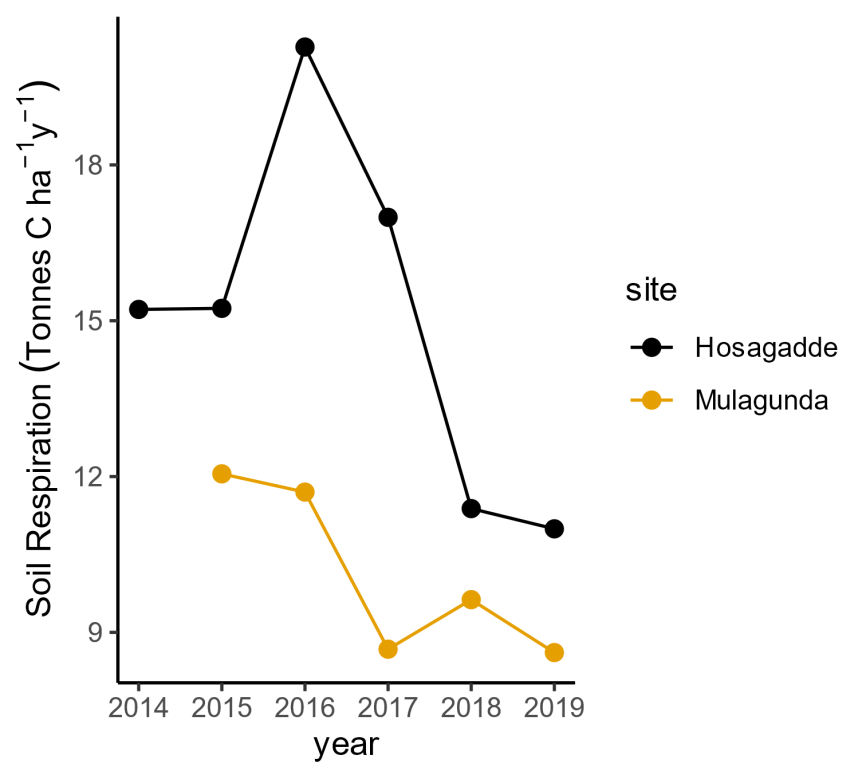
