## Supplemental Table 1 for "Soil respiration is correlated with rainfall and soil moisture at multiple temporal scales in a seasonal wet tropical forest"

*Table S 1: Summary of linear model results of Annual soil respiration to A) Annual rainfall*

*B) Mean annual temperature, C) Number of rainy days/year and D) Number of days in a year that have received a daily rainfall >50mm (referred to as heavy rainy days). Significant p-values are in bold.*

*Model A: Annual Rs (Mg ha^-1^) ~ Annual rainfall +site*

*Model B: Annual Rs (Mg ha^-1^) ~ Mean annual temperature +site*

*Model C: Annual Rs (Mg ha^-1^) ~ Annual number of rainy days +site*

*Model D: Annual Rs (Mg ha^-1^) ~ Annual number of heavy rainy days +site*

| **Model** | **Annual Rainfall +**  **Site** | | **Mean annual temperature +**  **Site** | | **No. Rainy days/year +**  **Site** | | **No. heavy rainy days/year +**  **Site** | |
| --- | --- | --- | --- | --- | --- | --- | --- | --- |
| *Predictors* | *Estimates* | *p* | *Estimates* | *p* | *Estimates* | *p* | *Estimates* | *p* |
| (Intercept) | 22.1257 | **<0.001** | 6.6343 | 0.873 | 0.8014 | 0.956 | 19.4290 | **<0.001** |
| Annual rainfall (mm) | -0.0017 | **0.018** |  |  |  |  |  |  |
| Site | -3.6821 | **0.023** | -5.1882 | 0.059 | -6.0053 | **0.019** | -3.9258 | **0.021** |
| Mean annual temperature (°C) |  |  | 0.3798 | 0.841 |  |  |  |  |
| Number of rainy days/year |  |  |  |  | 0.0872 | 0.344 |  |  |
| Number of heavy rainy days/year |  |  |  |  |  |  | -0.1565 | **0.030** |
| Observations | 11 | | 11 | | 11 | | 11 | |
| R^2^ / R^2^ adjusted | 0.750 / 0.687 | | 0.478 / 0.348 | | 0.534 / 0.418 | | 0.719 / 0.648 | |
